# Performance comparison and in-silico harmonisation of commercial platforms for DNA methylome analysis by targeted bisulfite sequencing

**DOI:** 10.1101/2021.03.12.435105

**Authors:** Miljana Tanić, Ismail Moghul, Simon Rodney, Pawan Dhami, Heli Vaikkinen, John Ambrose, James Barrett, Andrew Feber, Stephan Beck

**Affiliations:** University College London, UCL Cancer Institute, London, WC1E 6BT, United Kingdom; Institute for Oncology and Radiology of Serbia, Experimental Oncology Department, Belgrade, 11000, Serbia; University College London, Genomics and Genome Engineering Translational Technology Platform, London, WC1E 6BT, United Kingdom; University College London, Bill Lyons Informatics Centre, London, WC1E 6BT, United Kingdom; University College London, Division of Surgery and Interventional Science, London, W1W 7TS, UK; Royal Marsden Hospital, Molecular Pathology, London, SM2 5NG

**Keywords:** DNA methylation, technology comparison, epigenetics, benchmarking, targeted bisulfite sequencing, hybridization capture bisulfite sequencing, RRBS, genome-wide DNA methylation, epigenomics

## Abstract

DNA methylation is a key epigenetic modification in the regulation of cell fate and differentiation, and its analysis is gaining increasing importance in both basic and clinical research. Targeted Bisulfite Sequencing (TBS) has become the method of choice for the cost-effective, targeted analysis of the human methylome at base-pair resolution. Here we benchmarked five commercially available TBS platforms, including three hybridization capture-based (Agilent, Roche, and Illumina) and two RRBS-based (Diagenode and NuGen), across 16 samples. A subset of these were also compared to whole-genome DNA methylation sequencing with the Illumina and Oxford Nanopore platforms. We assessed performance with respect to workflow complexity, on/off-target performance, coverage, accuracy and reproducibility. We find all platforms able to produce usable data but major differences for some performance criteria, especially in the number and identity of the CpG sites covered, which affects the interoperability of datasets generated on these different platforms. To overcome this limitation, we used imputation and show that it improves the interoperability from an average of 10.35% (0.8M CpG sites) to 97% (7.6M CpG sites). Our study provides cross-validated guidance on which TBS platform to use for different features of the methylome and offers an imputation-based harmonization solution for improved interoperability between platforms, allowing comparative and integrative analysis.

## Introduction

DNA methylation is an indispensable epigenetic mark for many biological processes such as organismal development, cellular differentiation, and maintenance of cell-type-specific states^1^. Interrogating the changes in DNA methylation patterns is essential to better understand the biology of normal and pathological states and to identify clinically relevant biomarkers. In the past decade methods for methylome analysis have moved away from semi-quantitative methods with coarse resolution (MeDIP, MRE-seq) towards methods with the single-base resolution based on bisulfite conversion such as microarrays and next-generation sequencing (NGS)^2, 3^. NGS offers several advantages, including single-molecule analysis and read phasing to study sample heterogeneity and epiallele composition^4, 5^. Whole-genome bisulfite sequencing (WGBS) remains the gold standard method for studying DNA methylation at a single base-pair resolution, although it is associated with high costs and requires substantial computational resources. Targeted bisulfite sequencing (TBS) directs the sequencing to more informative parts of the genome, reducing the costs of sequencing. The analysis of DNA methylome by TBS can be achieved either through target-specific enrichment of regions containing CpG sites of interest using probe hybridization capture methods (HC)^6^ or through non-specific enrichment of CpG-dense regions by reduced-representation bisulfite sequencing (RRBS) mediated by a restriction enzyme recognition site containing a CG motif^7^.

Currently, five commercial manufacturers are producing off-the-shelf kits, offering standardized reagents and conditions for genome-wide TBS using HC (Agilent SureSelect Methyl-seq, Roche NimbleGen SeqCap EpiGiant, and Illumina TruSeq Methyl Capture EPIC) or RRBS (Diagenode Premium RRBS and Tecan NuGen Ovation RRBS Methyl Seq). The five platforms employ different experimental strategies to generate sequencing libraries and differ in the scope of regions they cover. The diversity of these platforms’ characteristics, along with the absence of a thorough comparison^8^ of their output and performance, warrants a comprehensive benchmarking to provide guidance for users to select the most appropriate platform based on each one’s strengths and limitations.

Here, we systematically compare these five platforms across a set of 16 samples, generating a total of 80 TBS libraries. We compare performance in terms of sequencing output, target capture efficiency, genomic features coverage, CpG coverage similarity, intra-platform reproducibility, between-platform concordance, and differential methylation. Finally, we benchmark each of the five TBS platforms to gold-standard data generated using WGBS and Nanopore sequencing.

## Results

### DNA methylation data generation

DNA methylome profiles were generated on a set of 16 samples (**Fig. 1a**), including 11 biological replicates that consisted of: reference gDNA isolated from human peripheral blood cells (Ref.gDNA) at two different DNA inputs (recommended by the manufacturer and 500 ng) in duplicate; Coriell NA12878 and Hela cell lines processed in duplicate; four DNA methylation standards generated from ZYMO fully methylated and unmethylated control samples; a pair of genetically and phenotypically divergent bladder cancer cell lines and a pair of isogenic cell lines with different sensitivity to cisplatin treatment (**Supplementary Table 1**). The same DNA from each sample was processed on each platform. The length and complexity of the library prep protocols vary considerably between the methods (**Fig. 1b**). Libraries were sequenced on Illumina HiSeq2500 at 100bp PE to achieve an average of 30M uniquely mapped deduplicated read pairs, and data were processed using the same computational pipeline. Additionally, whole-genome DNA methylation data was generated using WGBS on Illumina sequencer (Ref.gDNA) and Oxford Nanopore platforms (Coriell NA12878).

**Table 1:** Overview of platform design differences.

| Manufacturer | Agilent | Roche | Illumina | Diagenode | NuGen |
| --- | --- | --- | --- | --- | --- |
| Platform | SureSelect MethylSeq | NimbleGen SeqCap EpiGiant | TruSeq Methyl Capture EPIC | RRBS Premium | Ovation RRBS Methyl Seq |
| **Enrichment method** | in solution hybridization<br>150-mer biotinylated cRNA baits | in solution hybridization<br>60-90mer biotinylated DNA bait | in solution hybridization<br>biotinylated DNA bait | CCGG restriction enzyme site enrichment | CCGG restriction enzyme site enrichment |
| **Targeted region** | 84Mb | 80.5 Mb | 107 MB | 95.3 Mb** | 95.3 Mb** |
| **Number of targeted CpG sites** | 3,153,816 | 2,806,466 | 3,346,505 | 3 - 5 million (4,070,782**) | 3 - 5 million (4,070,782**) |
| **Recommended DNA input** | 1 µg | 1 µg | 500 ng | 100 ng | 100 ng |
| **Fragmentation method** | Ultrasonication | Ultrasonication | Ultrasonication | Restriction enzyme digestion (*MSpl*) | Restriction enzyme digestion (*MSpl*) |
| **Time-to-result\*** | 3 days | 5 days | 2 days | 3-5 days | 1 day |
| **PCR cycles** | 14 | 12 pre + 14 post-hybridization | 11 | 6-12 (determined by qPCR for each sample) | 12 |
| **Multiplexing** | 16 - 96 indexes | 24 indexes | 24 indexes | 24 indexes | 16 – 96 indexes |
| **Advantages** | Defined capture regions | Defined capture regions<br>Targets both strands<br>Post-BC-capture | Defined capture regions<br>Sample pooling -<br>Lower per-sample DNA input requirements | Sample pooling - reduced cost per sample and handling<br>Targets both strands | Short protocol<br>Unique molecular identifiers - precise duplicate removal<br>Targets both strands |
| **Disadvantages** | High per-sample DNA input requirements<br>Single strand capture | High per-sample DNA input requirements<br>Long protocol | Sample pooling - reduced flexibility for sequencing<br>Single strand capture | Non-specific CpG enrichment<br>Sample pooling - reduced flexibility for sequencing<br>No duplicate removal | Non-specific CpG enrichment |
| **Cost per sample\*\*\*** | \$\$ | \$\$ - \$\$\$ | \$\$ | \$ | \$ |

\*per user manual; \*\* *in silico* estimate; \*\*\* Cost per sample: \$ - <100\$; \$\$ - 100-500\$; \$\$\$ - >500\$

**Figure 1.**
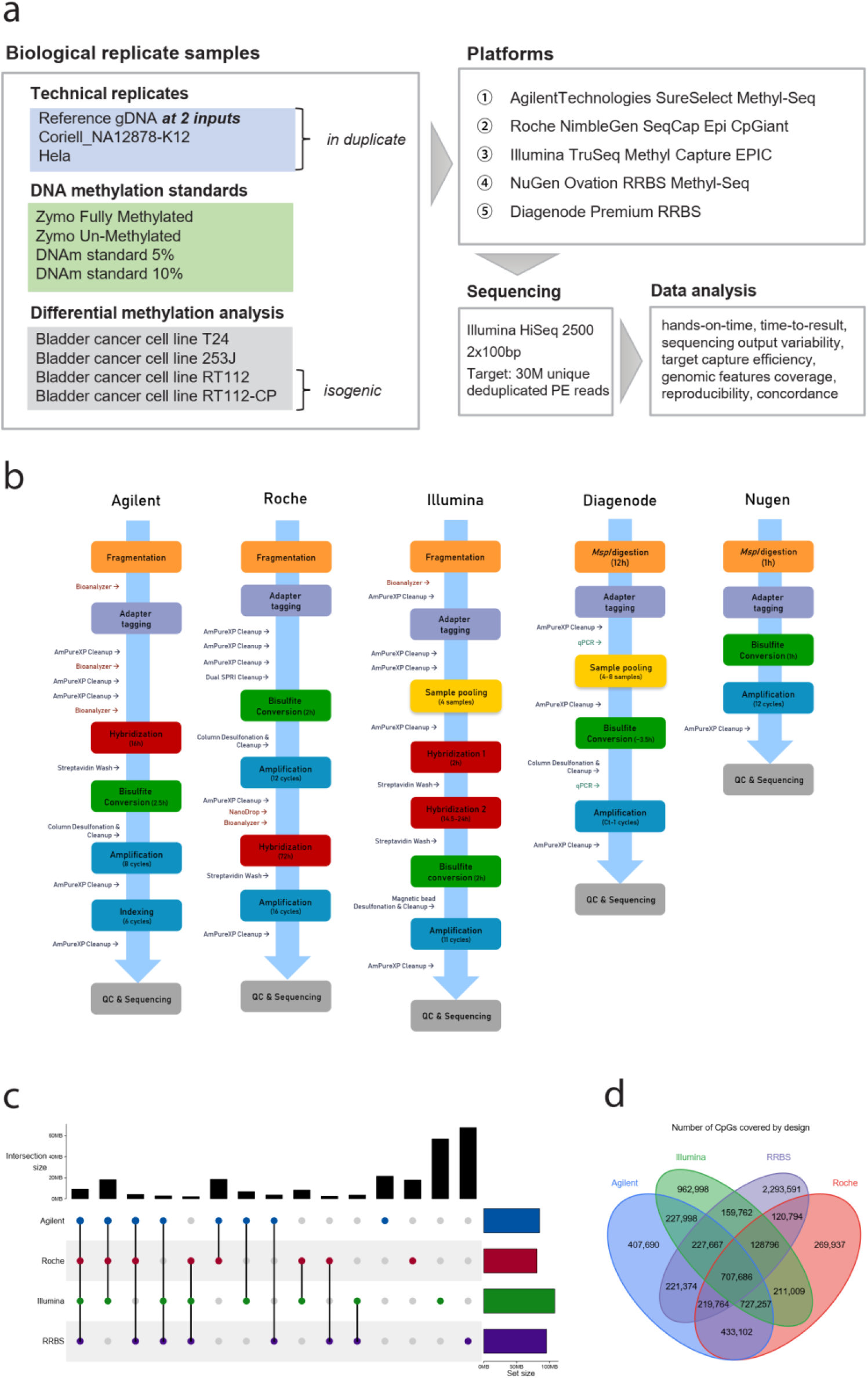
Technology and design comparison of targeted bisulfite sequencing platforms. (**a**) Schematic outline of the study design. **(b)** Schematic overview of the library preparation protocol steps for each of the platforms. **(c)** UpSet plot showing the overlap of regions targeted by design for Agilent, Roche, and Illumina hybridization capture platforms and *in silico* predictions for RRBS. **(d)** Venn diagram showing the overlap of CpG sites targeted by design for Agilent, Roche, and Illumina hybridization capture platforms and *in silico* predictions for RRBS.

### Differences in platform design

The five platforms differ in the methods used for DNA fragmentation, type of hybridization probes, the total size of the epigenome covered, the number of targeted CpGs, strand specificity, required DNA input, protocol complexity, total hands-on time and time-to-result, and finally the price per sample (**Table 1)**. While HC platforms target specific regions of the genome by design, RRBS non-specifically enriches for CpG-rich sequences.

To compare the overlap between the size of the regions covered by each platform manufacturers’ design files were used for the three HC platforms. To determine the theoretical target regions for RRBS platforms we performed an *in silico* enzymatic digestion with size selection mimicking experimental and sequencing conditions. All platforms commonly target only 9,260,583 bp, and 18,204,312 bp are shared by all HC platforms (**Fig. 1c)**. Agilent and Roche platforms commonly target additional 18,558,613 bp, while Illumina and *in silico* RRBS target the most platform-specific regions, 56,996,695 bp and 67,747,965 bp respectively. The size of target regions determines the amount of sequencing required but does not necessarily correlate with the number of enclosed CpG sites. The number of CpGs targeted by each platform ranged from 2.8 – 4 million CpGs. A total of 707,686 CpGs are commonly targeted by all platforms (**Fig.1d**). The three HC platforms commonly target 727,257 CpGs, and RRBS targets the highest number of platform-specific CpGs (2,293,591).

### Sequencing output variability

Cost-efficiency of sequencing depends on several factors including read-quality, mapping rates, duplication rates, and in case of hybridization capture methods, target capture efficiency. To simulate real-life biological experimental setup and to enable an unbiased comparison between platforms, equimolar amounts of each sample library were pooled together and sequenced to achieve an average of 30M of uniquely mapped deduplicated reads (UMDR). To achieve the targeted number of UMDR, between ∼55M reads (for NuGen) to over ∼80M reads (for Illumina’s platform) had to be sequenced (**Fig. 2a**). To determine the technical variability of sequencing output, we report the average number of reads and the standard deviation for each of the platforms tested (**Supplementary Table 2**). Due to reduced sequence diversity of bisulfite converted DNA, alignment rates are generally lower than for genome sequencing. Mapping rates for HC-based methods were comparable between platforms (Agilent: 76.5% ± 1.4%, Roche: 78.6% ± 3.9% and Illumina: 82.1% ± 1.8%) and higher than for RRBS-based platforms (Diagenode: 60.6% ± 1.4% and NuGen: 68.0 ± 3.4%) (**Supplementary Fig. 2**). Bisulfite conversion is a harsh chemical process that degrades DNA, reducing the library complexity and if combined with a high number of PCR cycles, may lead to a large number of duplicate reads skewing the methylation readout^9, 10^. Deduplication is required to determine precise CpG methylation levels and can be achieved by relying on fragment genomic coordinates^11^ (Agilent, Roche, and Illumina) or using unique molecular identifiers (UMIs)^12^ (Nugen). Duplication rates were the highest for Illumina’s platform (52.0% ± 16.4%) and the lowest for NuGen (7.0% ± 6.2%). Overall, the highest ratio of UMDR to passing-filter (PF) reads was observed for NuGen’s platform and the lowest for Illumina.

**Figure 2.**
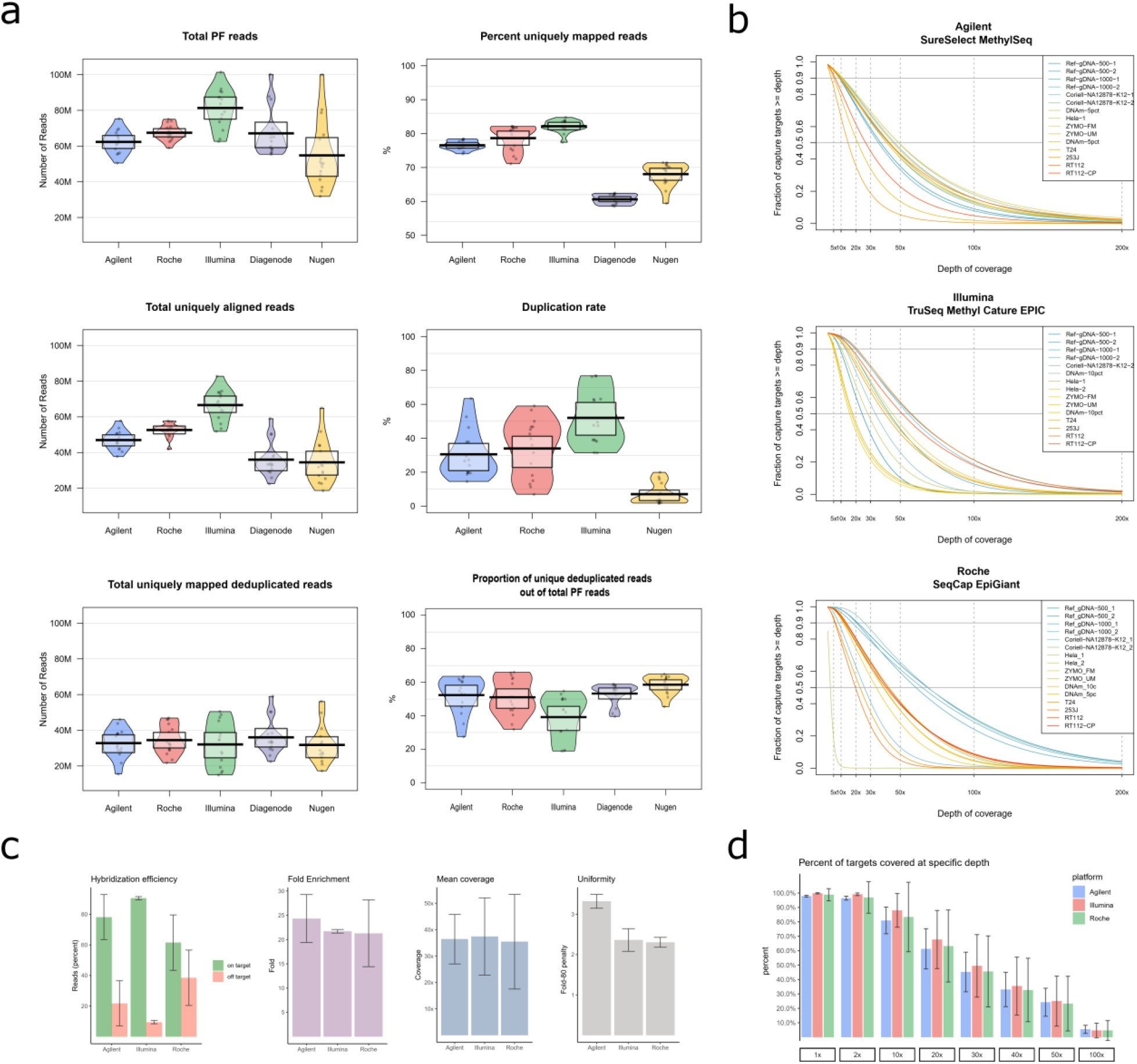
Sequencing performance by the platform. **(a)** Variability in sequencing output by each platform, showing the total number of passing filter (PF) reads (top left), alignment rate (top right), the total number of uniquely mapped PF reads (middle left), percent duplicated reads (middle right), the total number of deduplicated uniquely mapped PF reads (bottom left), and the efficiency of sequencing in terms of the proportion of usable reads (deduplicated uniquely mapped PF reads) over the total number of PF reads. **(b)** The fraction of targets covered at specific dept of sequencing for each sample by the platform. **(c)** Probe hybridization target capture performance measured as the percentage of on+near bait bases that are on as opposed to near, off-target as the percentage of aligned PF bases that mapped neither on nor near a bait, fold enrichment by which the baited region has been amplified above genomic background, the mean coverage of targets and the uniformity as the fold over-coverage necessary to raise 80% of target bases to the mean coverage level in those targets. **(d)** On-target target capture efficiency variability by platform represented as the mean +/- standard deviation of the fraction of targets covered at a specific depth.

### Target capture efficiency

Sequencing cost-effectiveness for HC methods also depends on the percentage of sequencing directed to regions of interest versus the amount of off-target sequencing. For this analysis, we omitted two RRBS-based platforms since their enrichment is not target-specific. In order to evaluate the performance of the wet-lab hybridization capture protocol for the three HC platforms (Agilent, Roche, and Illumina), on-target capture efficiency was compared based on coverage of all targeted bases according to manufacturers’ design files. Overall, on-target capture efficiency measured as the percentage of uniquely aligned PF bases that mapped on or near a bait, was highest for Illumina’s platform (90.6% ± 1%), followed by Agilent (78.2% ± 14.8%) and Roche (61.5% ± 18.2%) platforms, with the two latter platforms marked by high inter-sample variations (**Fig.2c** and **Supplementary Table 3**). Fold enrichment by which the baited region has been amplified above the genomic background was highest for Agilent (24.3 ± 7.2), followed by Illumina (21.7 ± 0.4) and Roche (21.3 ± 6.9).

Target coverage was assessed as a proxy for how well the data is likely to perform in downstream applications. Median target coverage was similar for all platforms, Illumina 31.5x ± 12.5x, Roche 30.3x ± 14.5x, and Agilent 27.4x ± 7.2x. The uniformity of coverage of the target region was represented as the theoretical increase of non-zero read coverage necessary to raise 80% of bases to the mean coverage levels in those targets (fold-80 penalty), with lower values indicating less variability in target coverage. We observed higher uniformity of coverage for Roche and Illumina’s platforms (2.3 ± 0.1 and 2.4 ± 0.3, respectively), while Agilent had a fold-80 penalty score of 3.3 ± 0.2. The uniformity of coverage translates into the fraction of targeted regions being covered at a specific depth (**Fig.2 b**). Intra-platform variability was pronounced for all three HC platforms, including technical replicates, with the average percentage of targets covered at 10x ranging from 80.9% ± 9.3% for Agilent’s platform to 88.1% ± 11.7% for Illumina (**Fig.2 d**).

### CpG coverage

An important consideration when choosing a platform for methylome analysis is the number of CpG sites that it interrogates. The number of CpGs covered by RRBS is a function of the depth of sequencing and sequencing conditions (read length and single- or paired-end mode). While each 40 – 600 bp fragment generated by restriction enzyme *MspI* has a CpG site at its ends, the number of CpGs within each fragment is not uniform due to the random selection of recognition site by the enzyme. In contrast to HC methods, the number of CpGs covered is determined by the target region. However, we observed a much higher number of experimentally covered CpGs at 1x than expected by design for all three HC platforms, consistent with high levels of off-target sequencing (**Fig. 3a**). The average number of CpG sites covered at least 5x drops to approximately 4 million CpGs for all platforms but remains higher for RRBS platforms at 10x and 30x compared to HC methods. Both RRBS platforms showed a similar number of CpGs covered at a specific depth, ranging from over 5 M CpGs covered at 1x, to ∼2 M CpGs covered at 30x. Mean CpG coverage was higher for RRBS methods (Diagenode: 33 ± 10.6, NuGen: 31 ± 9.6) compared to HC methods (Agilent: 13 ± 6, Roche: 9.7 ± 4.1 and Illumina: 13.2 ± 3.6), reflecting a more uniform coverage for the former.

**Figure 3.**
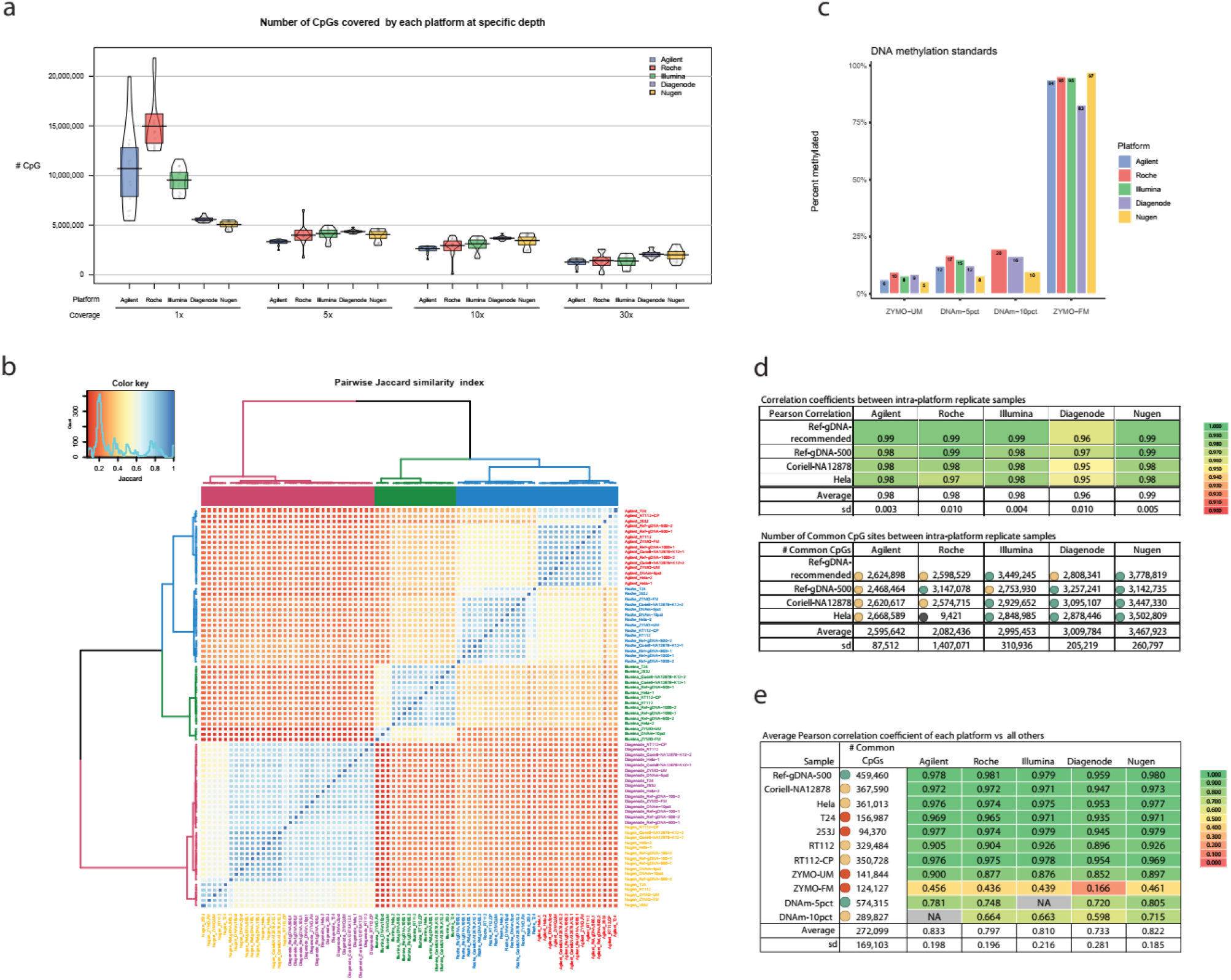
Platform reproducibility and concordance of DNA methylation calls. **(a)** Total number of CpGs covered at 1x, 5x, 10x, and 30x for each sample stratified by the platform. **(b)** The platform-sample similarity in terms of the identity of CpGs covered at 10x is shown as unsupervised hierarchical clustering of the pairwise Jaccard similarity index, representing the number of common CpGs over the union of CpGs. **(c)** Global methylation levels for DNA methylation standard control samples stratified by the platform. **(d)** Intra-platform reproducibility of DNA methylation levels per CpG. The top panel shows the pairwise Pearson correlation coefficient for duplicate samples per platform. The bottom panel shows the number of common CpGs covered at 10x for a given sample pair. **(e)** Inter-platform concordance in CpG methylation levels per sample. The average Pearson correlation coefficient of pairwise comparisons across all platforms for each sample (one platform *vs* other platforms), and the number of common CpGs covered at 10x for a given sample across all platforms compared.

Next, we wanted to determine the degree of CpG coverage similarity between and within platforms, in terms of the number of overlapping CpGs covered with a minimum of 10 reads. For each sample-platform, we calculated a pairwise Jaccard similarity index defined as the number of commonly interrogated CpG sites (intersection) over the total number of CpGs (union). Unsupervised hierarchical clustering separated samples into two main groups, RRBS-based and HC-capture-based, indicating that by large, they cover different CpG sites (**Fig. 3b**). All samples clustered according to the platform of origin, with Diagenode showing the highest degree of intra-platform similarity. As expected, NuGen and Diagenode RRBS platforms covered mostly the same CpG sites, while among HC methods, Agilent and Roche platforms had a higher degree of similarity compared to Illumina.

### Theoretical and empirical coverage of genomic features

The CpG distribution is not uniform throughout the genome; thus an important consideration for choosing a platform is the coverage of genomic features of interest. HC platforms target CpG islands (CGI), shores and shelves, regulatory regions, and known DMRs to a different extent by design. We compared the size of targeted genomic features and the number of CpGs targeted in each platform (**Supplementary Fig. 3**). The Illumina platform targets ∼20 Mb more than other platforms including the highest portion of FANTOM5 defined enhancers (>40%) and insulator regions. Agilent’s platform targets the largest region and the highest number of CpGs within CGI, in promoter end exon genic regions. Interestingly, although Illumina’s platform targets the largest portion of inter-CGI, intron, and heterochromatin regions, RRBS encompasses more CpGs in those regions.

To determine how well each platform experimentally covers specific regions of the genome, we counted the number of CpGs with a minimum of 10 reads per platform in each genomic/regulatory feature (**Fig. 4a**). In general, we observed a pronounced variation in the number of CpGs covered per sample by feature for all platforms except Diagenode. Striking differences in theoretical vs empirical coverage were observed, especially for Agilent’s platform reflecting lower uniformity of coverage with many CpGs not reaching the 10x threshold. Overall, RRBS platforms covered the highest number of CpGs located in CGI, shores, and shelves were best covered by Roche, while open-sea CpGs were best represented by Agilent and Roche. Illumina interrogated over twice as many CpGs located in FANTOM5 enhancer regions compared to other platforms. Roche showed the highest coverage of promoters, exons, 5’ and 3’UTRs, while RRBS platforms targeted most CpGs in introns. When looking at regulatory features defined by ChromHMM^13^ where promoters were stratified by their activity, HC methods covered more CpGs in active promoter regions, while RRBS methods had higher coverage of weak promoters. There was no significant difference between platforms in the coverage of either strong or weak enhancers, whereas Illumina covered the most CpGs located in insulator regions.

**Figure 4.**
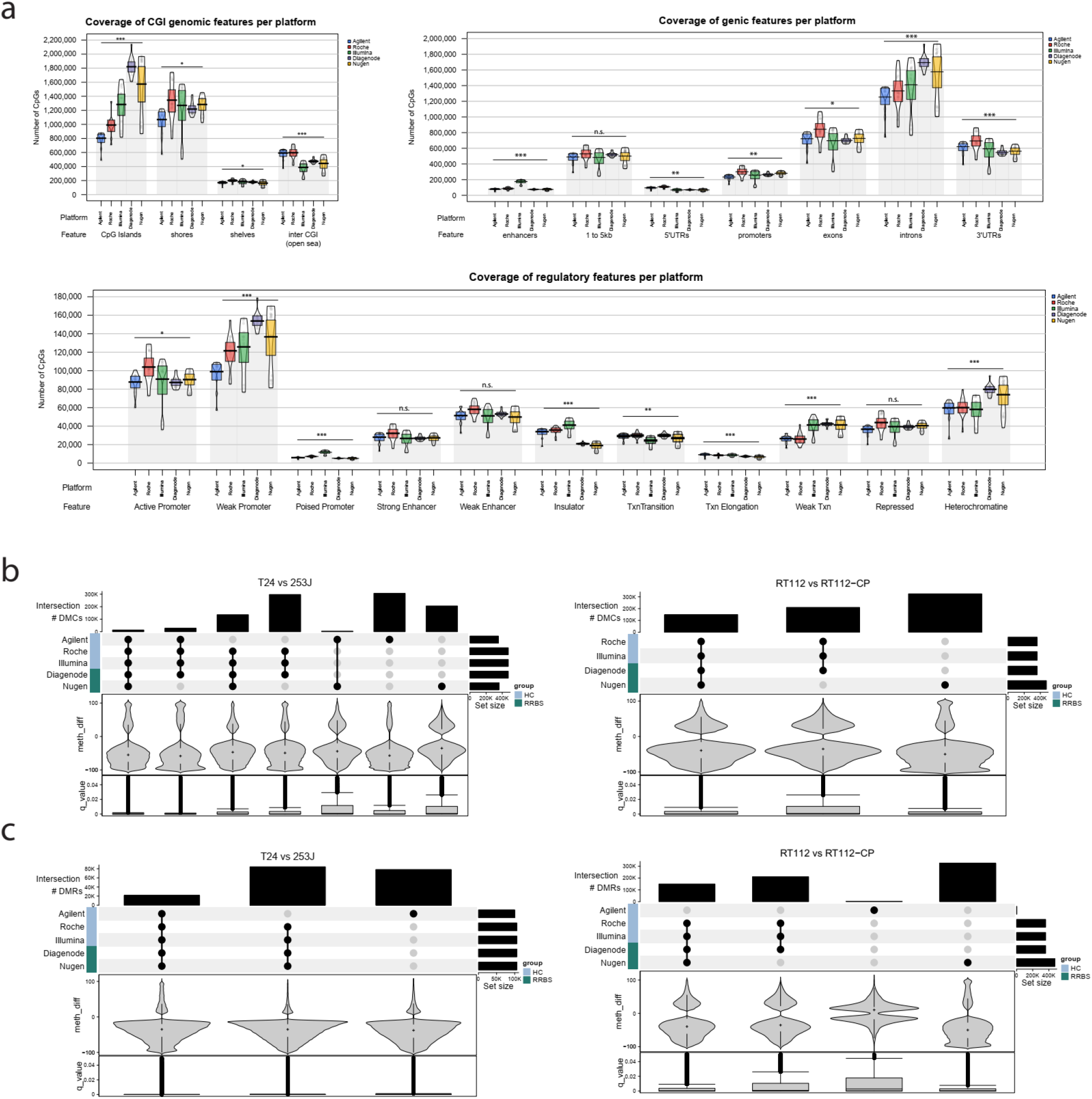
Feature annotation and differential methylation calls by the platform. **(a)** Experimental variability in the number of CpGs covered at 10x stratified by genic features (top left), CpG density (top right), and regulatory features (bottom) for each platform. **(b)** UpSet plot showing the overlap of differentially methylated CpG sites (DMC) called by each platform in genetically distinct pairs of cell lines (left) and pair of isogenic cell lines (right). **(c)** UpSet plot showing the overlap of differentially methylated regions (DMR) called by each platform in genetically distinct pairs of cell lines (left) and pairs of isogenic cell lines (right).

### Intra-platform reproducibility and inter-platform concordance

To estimate the level of confidence in obtained results for any given biological experiment and to set appropriate cutoffs for calling differential DNA methylation, it is important to know the approximate level of technical variation in the data. To measure the contribution of technical variation and to assess platform reproducibility we compared DNA methylation data from samples for which the same DNA was used to generate duplicate sequencing libraries (Ref.gDNA, Coriell, and Hela). DNA methylation levels were calculated as the fraction of methylated CpGs over the sum of methylated and unmethylated CpGs. We constricted the analysis to CpG sites covered at a minimum of 10x in both samples to calculate pairwise Pearson correlation coefficients. All platforms showed a very high overall correlation in DNA methylation levels of replicate samples, with Diagenode slightly underperforming (**Fig. 3d** and **Supplementary Fig. 5**). The effect of DNA input between platforms was estimated using Ref.gDNA at different inputs for library prep (**Supplementary Fig. 4**). HC methods require ∼10-fold higher amounts of DNA compared to RRBS. For HC methods, 1 ug of DNA (recommended for Agilent and Roche) was compared to 500 ng input (recommended for Illumina), while for RRBS methods recommended 100 ng input was compared to 500 ng (common to all platforms). All platforms, except for Diagenode, showed very high intra- and inter-platform correlation in DNA methylation levels (>0.98) for different DNA inputs.

Concordance between platforms for overall DNA methylation levels was compared on a set of DNA methylation standards generated by mixing DNA from ZYMO unmethylated and ZYMO fully methylated controls at know ratios (0% ZYMO-FM, 5% ZYMO-FM, 10% ZYMO-FM, 100% ZYMO-FM). ZYMO-UM control DNA is isolated from DNMT1 and DNMT3b knockout cell line known to have less than 5% methylated DNA^14^. All technologies appeared to slightly overestimate methylation levels for unmethylated control and to underestimate methylation levels of fully methylated control (**Fig. 3c**). Methylation levels for unmethylated control ranged from 5% estimated by NuGen to 10% estimated by Roche, while for the fully methylated control, levels ranged from 83% (Diagenode) to 97% (NuGen).

We next looked at the between-platform concordance in methylation levels for common CpG sites for each sample in a pairwise fashion. The intersection of CpGs covered by all five methods was made to enable the direct comparison. Methylation levels per sample for each platform were compared to all others to calculate the average correlation over common CpGs (**Fig. 3e**). Roche Hela-1 replicate was excluded from the analysis since the hybridization capture failed. The number of commonly covered CpGs in all samples ranged between 94,000 to 450,000 CpGs. We observed a high correlation in called methylation levels for all samples (R:0.85-0.99) except for DNAm-5pct, DNAm-10pct, and ZYMO fully methylated control (**Supplementary Fig. 6**).

### Imputation to improve selected CpG site concordance

The platform similarity as a function of the number of overlapping CpG sites is of particular concern as downstream analyses regularly require subsetting to a set of CpG sites that are present in all datasets. Furthermore, the effect of low overlap in the CpG sites selected is further exacerbated as the number of samples/datasets increases. Thus, samples with duplicate sequencing libraries (Ref.gDNA and Coriell) were further analyzed to measure the effect on the concordance in the CpG sites selected across platforms. A total of 7.8M CpG sites were covered at 10x by the five platforms, of which only 9.85% (0.79M CpGs) are present in all five platforms.

Imputation can be used to estimate the methylation value for each missing CpG site ^15, 16^ and thereby, can be used to help increase platform intra- and interoperability. We used BoostMe^16^ to impute the vast majority (i.e. 99.95%) of missing CpG sites, leveraging information learnt from just the neighboring CpG within the same dataset. The overlap between the five platforms increased almost tenfold (0.79M CpG sites to 7.6M CpG sites) (**Figure 5c**). However, it comes at a cost of reduced DNA methylation concordance from an average Person correlation of 0.97 before imputation to 0.80 after imputation (**Supplementary Table 5** and **Supplementary Figures 8-10**). Mean absolute error (MAE) that estimates by how much each methylation value is off, increased from an average 0.05 to 0.14, while MAE corrected for the number of CpGs present in the dataset, decreased from an average of 0.12 to 0.002. When a distance threshold of <25bp was applied to impute neighboring CpGs the total number of overlapping CpGs was 2.5 million (32% of the total), while the intraplatform concordance after imputation remained high (ρ=0.89 and MAE=0.10).

**Figure 5.**
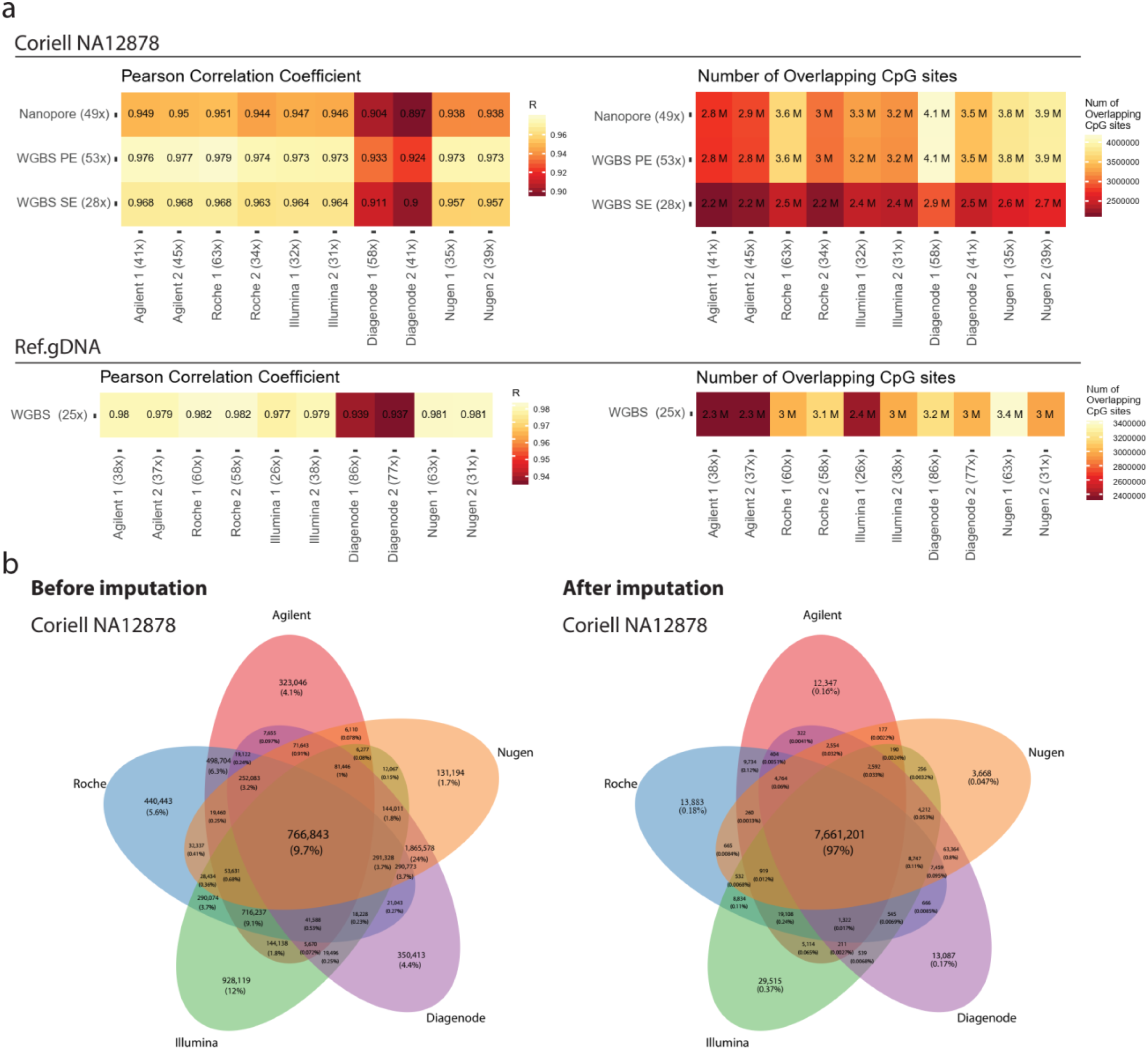
**a)** Pairwise comparisons between TBS platforms and Coriell NA12878 WGBS and Nanopore data (top) and Ref.gDNA WGBS data (bottom). Each tile in the left pane shows showing Pearson correlation coefficients and in the right pane refers to the number of overlapping CpG sites between the two samples compared. **b)** Number of overlapping CpGs for Coriell NA12878 before and after imputation with BoostMe.

### Calling differentially methylated sites and regions

Ultimately, many studies aim to identify differentially methylated CpG sites (DMC) or differentially methylated regions (DMR) between samples. To assess the between-platform concordance for calling differential methylation, we compared two pairs of bladder cancer cell lines, one pair with a different genetic origin and diverse phenotypes (253J and T24) and a second pair of isogenic cell lines with divergent sensitivity to cisplatin (RT112 and RT112-CP). Given that the observed technical variability was as high as 5% for some platforms, we set the threshold to over 10% methylation difference and q-values less than 0.05 to call DMCs and DMRs as differentially methylated. Only a small subset of DMCs was commonly identified by all five platforms for both pairs of cell lines (**Fig.4 b**). For the T24 vs 253J cell pair, Agilent and NuGen platforms identified smaller sets of DMCs, of which many were unique with somewhat higher q-values. Surprisingly, with Agilent’s platform data we failed to call any significant DMCs, although the other four platforms shared most of the identified significant DMCs. As expected, the bulk of identified DMCs for all platforms showed higher methylation differences for the genetically distinct pair of cell lines compared to the isogenic pair. When comparing aggregated methylation levels over 1000 kb tiling windows to call DMRs, high concordance was observed between all five platforms for T24 *vs*. 253J cells, with NuGen identifying an additional subset of highly significant DMRs. Again, Agilent’s platform didn’t identify any of the DMRs called by the other four platforms for the RT112 *vs*. RT112-CP pair. Notably, methylation differences for significant DMRs were lower compared to methylation differences observed for DMCs.

### TBS comparison to WGBS and Nanopore methylome

Whole-genome methods for DNA methylation analysis can theoretically capture all CpGs at single base-pair resolution. WGBS is considered the gold standard for near-complete characterization of DNA methylome, yet it is liable to bias induced by bisulfite treatment^17^. The sequencing of the native DNA molecule using third-generation sequencing technologies platform allows for direct detection of modified bases but requires large sample inputs, and features relatively high error rates^18, 19^. Both approaches are costly in terms of computational and financial resources and therefore not suited for large sample cohorts. We compared TBS to whole methylome datasets. All datasets were filtered to exclude CpG sites with less than ten reads or with extremely high coverage (>3 sd). Ref.gDNA was sequenced by WGBS covering 18,505,516 CpGs (25x). All TBS platforms showed high levels of concordance for all platforms (ρ>0.98) with Diagenode slightly underperforming (ρ=0.94). Coriell NA12878 was represented by two WGBS datasets and a Nanopore data: WGBS SE covering 24,074,147 CpGs at 28x mean coverage and WGBS PE with 25,093,057 CpGs at 53x, whilst Nanopore covered 28,230,385 CpGs at mean 49x coverage. Correlation in methylation levels between WGBS SE and PE datasets was higher (ρ=0.95) than with Nanopore (ρ=0.91 and 0.93, respectively) (**Supplementary Figure 7**). All TBS platforms displayed a higher correlation to WGBS datasets than to Nanopore data, with Diagenode’s platform showing the lowest correlation, albeit with the highest CpG overlap.

## Discussion

To answer important biological questions in the epigenetics of health and disease, we must understand the possibilities and limitations of the tools we use. Over the past decade, many studies harnessed the power of BS-seq methods to advance our knowledge on epigenetics of non-communicable diseases, to identify clinical biomarkers and understand normal development. Several companies offer off-the-shelf products for targeted analysis of DNA methylation at the genome-wide level. We compared five commercial platforms (Agilent SureSelect Methyl-seq, Roche NimbleGen SeqCap EpiGiant, and Illumina TruSeq Methyl Capture EPIC, Diagenode Premium RRBS, and Tecan NuGen Ovation RRBS Methyl Seq), and examined their performance, similarities and differences. To mimic a real-life experimental setup, the same panel of 16 samples was processed on each platform, sequenced to a comparable number of usable reads, and analyzed using the same bioinformatics pipeline.

Small-scale studies focused on in-vitro experiments have fewer constraints compared to high-throughput applications using scarce clinical samples that dictate the choice of the platform. The time-to-result, protocol complexity, and required sample input differ considerably between platforms. NuGen offers the shortest and least complex protocol that requires only 100 ng. Among HC platforms, Illumina’s protocol can be completed in two days and requires half the amount of DNA by multiplexing four samples, albeit reducing the flexibility in experimental design. All platforms can be potentially automated with modifications of the bisulfite conversion step. The price per sample (for library prep only) for RRBS platforms is almost tenfold lower compared to HC platforms.

While all platforms perform adaptor tagging through ligation before the bisulfite conversion step, Agilent and Illumina perform hybridization capture before bisulfite conversion reaction whereas Roche platform performs target capture after bisulfite conversion.^20^ To distinguish between C>T SNP and C>U conversion it is necessary to read both strands for a given CpG site. In addition to RRBS platforms, only Roche HC platform targets both strands at the same ratio, while Illumina covers both strands only at common SNPs.

Differences in design, size of the targeted region, and experimental strategies for library preparation and target capture translate into stark differences in the cost-effectiveness of sequencing. To achieve a similar number of uniquely mapped deduplicated reads, HC platforms required more sequencing than RRBS platforms. Although all platforms slightly underestimated global 5mC levels for fully methylated DNA standards, this was particularly pronounced for Diagenode’s platform. *Olova et al*. determined that the bisulfite conversion is the main driver of sequencing biases that are further amplified by PCR amplification.^17^ Estimates of 5-methylcytosine (5mC) levels may be biased by under- or over-conversion, and/or incomplete duplicate removal. Since the sample source and sequencing conditions were the same for all, the bias must have been introduced during the library preparation step. Read duplication levels were highest for Illumina’s and lowest for NuGen’s platform. While HC platforms processing included a deduplication step based on read coordinates to account for bias introduced by PCR amplification, only NuGens’s platform incorporates unique molecular identifiers enabling precise read deduplication. In contrast, Diagenode’s platform lacks any means for read deduplication, rendering it susceptible to biased estimation of 5mC levels. Conversion efficiency was above 99% for all platforms except for Roche, where the rate was over 98%. Overconversion may lead to underestimation of 5mC levels; however, fully methylated spike-in was not included in all platforms so we could not compare the over-conversion rates.

The choice of method defines the space for the discovery of CpG loci or regions associated with a specific phenotype. Each of the platforms targets different regions of the genome and the overlap between CpG loci covered by all is modest. Overall, RRBS methods covered more CpGs at 10x, but almost 40% of those CpGs fall into repetitive regions (data not shown), predominantly Alu elements, which are not targeted by HC platforms. We provide a breakdown of genomic feature coverage by platform that can be used as a guide for platform selection. A pronounced variation in the number of covered CpGs within each platform was observed warranting an imputation of missing data to fully capitalize on the dataset. To increase power for the discovery of novel biomarkers meta-analysis and integration of datasets generated using different platforms can be used. The interoperability of the platforms is dependent on the number of overlapping CpGs between platforms and their concordance. Our results reveal high between-platform concordance in measured 5mC levels of the overlapping CpG sites. We found that while many of the DMCs called by the five platforms do not overlap, the majority of DMRs are commonly identified by all platforms, suggesting that region-level analysis rater than per-CpG should be performed. Alternatively, imputation of missing CpG sites applying distance treshold is a viable option to incrase inter-platform operability for downstream analysis.

In summary, our study provides an in-depth analysis of the TBS platforms’ comparative performance and characteristics allowing users to determine the optimal technology for methylome analysis.

## Materials and Methods

### Sample origin and processing

The reference genomic DNA (gDNA) sample (Ref.gDNA) was obtained by pooling DNA extracted from peripheral whole blood cells (PWBC) of adult healthy volunteers, providing informed consent under 15/YH/0311 of the UCL BioBank for Health and Disease. Blood samples were collected in EDTA-treated 10 ml BD Vacutainer™ tubes and centrifuged at 1900x g for 10 minutes at 4°C using a refrigerated centrifuge to separate blood cells from plasma. Buffy coat (0.5 ml) was transferred using Pasteur blob to a clean Falcon tube containing 4.5 mL HEMAgene®•BUFFY COAT DNA stabilizing reagent (Oragene) and the DNA was extracted using QIAamp DNA Blood Mini Kit (Qiagen). The Coriell-NA12878 DNA sample was purchased from Coriell Cell Repositories. Hela cell genomic DNA was purchased from New England Biolabs. Bladder cancer cell lines were acquired from the ATCC and cultured in prescribed media, gDNA was extracted using QIAamp DNA Mini Kit (Qiagen). The Human Methylated & Non-methylated DNA Set (ZYMO Research) was used to generate DNA methylation standards: fully methylated (ZYMO-FM), 10% and 5% methylated (DNAm-10pct and DNAm-5pct) and unmethylated (ZYMO-UM). The negative control (ZYMO-UM) DNA in the set comes from a Human HCT116 DKO Non-methylated DNA that contains genetic knockouts of both DNA methyltransferases DNMT1 (-/-) and DNMT3b (-/-) and has low levels of DNA methylation^14,^ while the fully methylated DNA control was generated by the manufacturer by treating the HCT116 DKO with *M*.*SssI* methyltransferase. Sample quality was determined using Agilent 2100 Bioanalyzer 1000 DNA chip to assess the DNA integrity and sample purity was estimated using NanoDrop spectrophotometer. Quantification was performed using Qubit dsDNA BR Assay (Invitrogen).

### Library preparation and target enrichment with Agilent SureSelect MethylSeq kit

To generate SureSelect MethylSeq sequencing libraries (Agilent Technologies Inc.) we followed the manufacturer’s protocol. Briefly, we sheared 1 µg of the gDNA on Covaris S2 ultrasonicator system to obtain ∼150bp fragments Fragmented DNA was cleaned up using AMPure XP beads (Beckman Coulter) followed by end repair, A-tailing, second clean up and adapter ligation before the final clean up. Quality and quantity of the libraries were checked using Bioanalyzer dsDNA High Sensitivity chip (Agilent Technologies) and Qubit HS dsDNA Assay (Life Technologies, Thermo Fisher Scientific) before overnight hybridization (min 16h) with biotinylated oligo RNA baits followed by streptavidin-conjugated magnetic bead pulldown and wash steps. The captured library was then bisulfite converted using EZ DNA Methylation-Gold Kit (Zymo Research), followed by 8 cycles of PCR amplification and AMPure XP bead cleanup. Second PCR amplification with 6 cycles of the bisulfite-converted libraries was performed to introduce sample indices, followed by final AMPure XP bead cleanup. Library concentration was measured using the Qubit dsDNA HS kit and the library size distribution on a Bioanalyzer High Sensitivity DNA chip.

### Library preparation and target enrichment with Roche NimbleGen SeqCap Epi kit

NimbleGen SeqCap EpiGiant libraries (Roche Sequencing) were prepared according to manufacturer’s instructions, as follows: 1 µg of the gDNA sample was sheared using Covaris S2 ultrasonicator system to an average DNA fragment size of 200 bp, sheared DNA was end-repaired, cleaned up with magnetic beads, followed by A-tailing of the 3’end, second clean up, adapter ligation and double SPRI cleanup. Libraries were bisulfite converted using EZ DNA Methylation Lightning Kit (Zymo Research), followed by 12 cycles of Pre-Capture LM-PCR amplification using Kapa Hifi Uracil+ polymerase, cleanup, and fragment distribution analysis on Bioanalyzer High Sensitivity DNA chip. The bisulfite-converted library was hybridized with biotinylated DNA capture probes over 72h and washed with streptavidin-conjugated magnetic beads. The enriched libraries were amplified by a second round of LM-PCR with 16 cycles using Kapa Hifi Uracil+ polymerase (Roche) and purified with AMPure XP magnetic beads. Library concentration was measured using the Qubit dsDNA HS kit, purity using NanoDrop 2000 spectrophotometer (ThermoFisher Scientific), and the library size distribution on a Bioanalyzer High Sensitivity DNA chip.

### Library preparation and target enrichment with Illumina TruSeq Methyl Capture Epic kit

The TruSeq Methyl Capture Epic kit was a gift from Illumina. Sequencing libraries were prepared according to the manufacturer’s instructions. In brief, 500 ng of each gDNA samples were sheared on Covaris S2 ultrasonicator to a median size of 160 bp followed by a cleanup step using AMPure XP magnetic beads, end-repair reaction, second magnetic bead clean up, A-tailing, adaptor ligation, and final clean up. Four samples containing adaptors were pooled together to a total mass of 2 µg and hybridized with a biotinylated DNA capture panel for 35min at 58°C. Target DNA fragments were captured by streptavidin-conjugated magnetic beads and washed, followed by a second overnight hybridization (touchdown from 95°C to 58°C) streptavidin-conjugated magnetic bead pulldown and wash step. Enriched libraries were bisulfite converted using EZ DNA Methylation Lightning kit with magnetic bead desulphonation and clean up. Eleven cycles of PCR amplification were performed using Kapa Hifi Uracil+ polymerase available separately from the kit. Final libraries were bead purified to remove adapters, quantified using the Qubit dsDNA HS kit and the library size distribution was checked on a Bioanalyzer High Sensitivity DNA chip.

### RRBS library preparation using Diagenode’s RRBS Premium kit

The protocol for Premium RRBS kit (Diagenode) library preparation included enzymatic digestion of 100 ng of gDNA with *MspI*, followed by end repair, adaptor ligation, and size selection by bead purification. Each library was quantified by qPCR to determine library concentration and 8 samples were pooled in equimolar amounts determined by an Excel pooling tool provided by the manufacturer. Pooled libraries were bisulfite converted using manufacturer-provided BS Conversion Reagent and desulphonated on columns. The second qPCR was performed to determine the optimal number of cycles for the final amplification step for each pool (11 and 12 cycles) before final cleanup. Library concentration was measured using the Qubit dsDNA HS kit and the library size distribution on a Bioanalyzer High Sensitivity DNA chip.

### RRBS library preparation using NuGen’s Ovation RRBS Methyl-Seq System 1–16

The Ovation RRBS Methyl-Seq System 1–16 (NuGen, now Tecan) user manual was followed to generate sequencing libraries by enzymatically digesting 100 ng of gDNA using *MspI*, followed by end repair, adaptor ligation, and final repair step. Generated libraries were bisulfite conversion using EpiTect Fast DNA Bisulfte Kit (Qiagen) purchased separately from the kit. Converted libraries were amplified by PCR using 12 cycles and purified using Agencourt® RNAClean® XP magnetic beads, and fragment distribution was checked on a Bioanalyzer High Sensitivity DNA chip. RRBS design BED file was created by *in silico* cutting of hg19 sequence at CG within CCGG motif and filtering to >100bp length ranges and 100 bp window from each end.

### Sequencing on an Illumina HiSeq 2500 instrument

All generated targeted bisulfite-converted libraries were sequenced on the HiSeq 2500 using HiSeq v4 SBS reagents (Illumina) at 2×100 bp. Barcoded samples per each library were multiplexed and libraries were denatured with NaOH and diluted to a final concentration of 2nM. Denatured libraries were loaded onto cBot for cluster generation on separate flow cell lanes in HighOutput or RapidRun mode according to the manufacturer’s protocol. Loading concentrations and percent of PhiX spike in libraries for each platform are as follows: Agilent at 15 pM with 5% of equimolar PhiX, Roche 16.5 pM with 5% equimolar PhiX, Illumina at 11 pM with 10% equimolar PhiX, Diagenode at 16 pM with 5% of 12 pM PhiX, and NuGen at 11.5 pM with 5% of equimolar PhiX. Sample pools were sequenced over several lanes/runs and raw FASTQ reads were merged prior to the analysis.

### Whole-genome bisulfite sequencing

Whole-genome bisulfite sequencing (WGBS) libraries for Ref. gDNA samples (500 ng) were generated using CEGX TrueMethyl® Whole Genome Workflow (WGW) Kit (CEGX, UK) with an average insert size of 600bp. Libraries were sequenced on Illumina X Ten at 2×150 bp with 1000M PE reads per lane. Read 2 was of low quality reducing the mapping down to 35% in PE mode. Therefore, only read 1 was used for mapping with a 78% alignment rate. WGBS_SE data for the Coriell NA12878 sample was downloaded from GEO public repository under accession number GSE86765 (Encode Project - ENCBS232JAP, 2x 125bp), libraries generated with were sequenced on Illumina HiSeq 2500, and WGBS_PE data from Illumina BaseSpace Hub MethylSeq - 8 replicates of Coriell NA12878 were prepared with the TruSeq Methylation kit, run on one flowcell of the HiSeq 4000 at 2×76. Replicates were combined across lanes to obtain 2- and 3-lane builds.

### Nanopore sequencing

One microgram of genomic DNA from the lymphoblastoid Coriell-NA12878 cell line was sheared using g-Tube (Covaris) to obtain high molecular weight fragments ranging from 50kb to 160 kb. Fragment ends were repaired using NEBNext End Prep kit (NEB) and AMX adapters were introduced using LSK109 ligation sequencing kit (Oxford Nanopore). Approximately 150 GB of data was generated using 2 flow cells on PromethION.

### Raw sequencing data processing

For all targeted BS-seq platforms Bismark^21^ data analysis pipeline was applied with few platform-specific modifications. Samples were demultiplexed either onboard Illumina HiSeq2500 or offline using bcl2fastq Conversion Software v1.8.4 (Illumina). The quality of raw reads in FASTQ format was checked using FASTQC^22^ (**Supplementary Fig. 1**).

For each sample-platform FASTQ files from different sequencing, runs were merged before alignment. NuGen FASTQ reads contain additional 1-3 inserted nucleotides and were trimmed using a diversity trimming python script provided by the manufacturer^22^. Any contaminating adaptor sequence was removed using TrimGalore^23^ in paired-end mode with default conditions for adaptors and --*trim1* option for Agilent, Roche, Illumina while for Diagenode’s platform *--rrbs* option was used, and for NuGen read 2 adaptor was specified. Bismark v0.10.1^21^ was used for alignment to the human genome build hg19 in paired-end mode using Bowtie 2 and default settings. After alignment PCR duplicates in Agilent, Roche and Illumina were marked and discarded using *deduplicate_bismark* perl script. For RRBS data this form of deduplication is not recommended as all reads start with the same sequence. However, the NuGen’s platform contains Unique Molecular Identifiers (UMIs) that enable for precise removal of duplicates using Duplicate Marking python script provided by the manufacturer^22^. Bismark’s module *bismark_methylation extractor* with options *--ignore_r2 2, --no_overlap, --comprehensive, --merge_nonCpG, --bedGraph, -- gzip* was used to generate output files with methylation status for each individual CpG, coverage and genomic coordinates. Bismark’s *coverage2citosine* tool with and without *--merge_CpG* option was used to obtain methylation status for each CpG in the genome both strands along with trinucleotide context, strand, coverage, and methylation percent. The efficiency of bisulfite conversion was verified by the ratio of C to T conversion of CHG and CHH (non-CG) dinucleotides. Nanopore data was processed using *nanoposlish*^18^. MultiQC was used for visualizing aggregated quality metrics.

#### Data analysis

Downstream analysis was performed using Unix and R environment. We didn’t use Repeat masking for any of the analyses. For hybridization capture methods PICARD^24^ CollectHsMetrics tool v2.3.0 was used to obtain statistics on HC efficiency. BEDTools^25^ and custom sed and awk scripts were used to calculate and visualize the target region coverage as a function of read depth. To compare target breadth of coverage UpSet plot from *ComplexHeatmap*^26^ and Venn diagram from *ChipPeakAnno* R packages were used to visualize the intersection of genomic ranges and CpG sites.

Bismark produced strand-specific methylation calls were imported into *MethylKit* R package for downstream analysis including CpGs with more than 10 reads. To get per CpGs methylation levels independent on a strand, counts on both strands of a CpG dinucleotide were merged. To perform pairwise comparisons in methylation levels, samples of interest were merged into one object containing common CpGs. Pearson correlation was used to assess concordance in DNA methylation levels in a pairwise comparison matrix. Differential methylation analysis was performed using Fisher’s exact test or logistic regression to calculate P-values per CpG and over 1000kb tiling windows, corrected by SLIM method^27^ to obtain q-values. Differentially methylated regions/CpGs were selected based on q-value<0.05 and percent methylation difference larger than 10%.

Feature annotation was done using *anotatr* R package. The CpG islands for hg19 were retrieved through *AnnotationHub* package and CpG shores were defined as 2Kb upstream/downstream from the ends of the CpG islands, less the CpG islands, CpG shelves as another 2Kb upstream/downstream of the farthest upstream/downstream limits of the CpG shores, less the CpG islands, and CpG shores. The remaining genomic regions make up the inter-CGI annotation. The genic annotations are determined by functions from *GenomicFeatures* and data from the *TxDb* and *org*.*hg19*.*eg*.*db* packages. Genic annotations include 1-5Kb upstream of the TSS, the promoter (< 1Kb upstream of the TSS), 5’UTR, first exons, exons, introns, CDS, 3’UTR, and intergenic regions (the intergenic regions exclude the previous list of annotations). FANTOM5 permissive enhancers determined from bi-directional CAGE transcription (Andersson et al. (2014) were added to genic annotations. Chromatin states were determined by hidden Markov models (chromHMM) on ENCODE histone mark data for nine cell lines: Gm12878, H1hesc, Hepg2, Hmec, Hsmm, Huvec, K562, Nhek, and Nhlf, were used to check the theoretical coverage of regulatory features by each platform ^13^. GRangesList object containing all cell lines data was stratified by chromatin state. Since cells have different regulatory states, overlapping ranges within each chromatin state were reduced ignoring strand so that each state in the plot represents the union of ranges for all cell types.

To calculate the breadth of coverage for the design regions determined based on manufacturer-provided BED files with annotation tracks, overlapping region pairs were intersected ignoring strand. To count the number of CpGs in regions of interest, the corresponding hg19 sequence was retrieved from BSgenome.Hsapiens.UCSC.hg19 using DNA strings R package, reduced to contain only non-overlapping regions independent of DNA strand and number of occurrences of CG was counted. The number of CpGs covered at a specific depth per platform was calculated using a custom R script. Jaccard similarity index was calculated using functions from *HelloRanges* R package. *Intervene* shiny app was used to visualize the Jaccard similarity index matrix^28^.

### Imputation

We used BoostMe^16^ to impute the vast majority (i.e. 99.95%) of missing CpG sites, leveraging information learnt from just the neighboring CpG within the same dataset. BoostMe was run with a 60:20:20 split between the training, validation and test datasets. On average, the training dataset was formed from 2M randomly selected CpG sites, while the validation and test datasets were made up of 0.67M randomly selected CpG sites. Imputation accuracy was 92.1% and 0.971 AUROC in cross-validation analysis, and 92.1% accuracy and 0.971 AUROC in the test dataset. In Imputation can be difficult in allosomes as the methylation composition can be significantly different from autosomal chromosomes due to the samples’ sex. As such, after excluding the allosomes, there is an average of 7.58M CpG sites, of which, an average of 4.23M CpG sites are missing in each dataset. BoostMe achieves a 92.1% accuracy and 0.971 AUROC in cross-validation analysis and achieves 92.1% accuracy and 0.971 AUROC with the test dataset (i.e., the dataset not used for training or cross-validation). In this manner, by imputing missing CpG sites with BoostMe, the overlap between the five platforms increases from 10.35% (0.79M CpG sites) to 97% (7.6M CpG sites).

## Declarations

## Acknowledgements

MT received funding from the European Union’s Seventh Framework Programme (Marie Skłodowska-Curie Actions FP7/2007-2013/WHRI-ACADEMY-608765); the Danish Council for Strategic Research (1309-00006B), the Ministry of Education, Science and Technological Development of Serbia (2011-2019/III-41026 and 451-03-68/2020-14/200043), and the Science Fund of the Republic of Serbia (PROMIS/2020/6060876). IM is supported by the Biotechnology and Biological Sciences Research Council (Grant No. BB/M009513/1). SB has received funding from the Wellcome Trust (218274/Z/19/Z) and a Royal Society Wolfson Research Merit Award (WM100023). AF received support from UCL/UCLH Biomedical Research Centre, Medical Research Council (MR/M025411/1), Prostate Cancer UK (MA_TR15_009) and BBSRC (BB/R009295/1). SR received funding from Orchid. We further acknowledge support from Dr Daniel Turner and Dr Botond Sipos (Oxford Nanopore Technologies) for the generation of the Nanopore-seq data and from the CRUK–UCL Centre-funded Genomics and Genome Engineering and Bioinformatics Translational Technology Platforms.

## Authors contributions

M.T., A.F. and S.B. conceived and designed the study. M.T. and S.R., performed the hybridization capture and RRBS experiments. P.D. and H.V. sequenced the libraries. M.T., and J.B. processed raw sequencing data. M.T. performed data analysis. I.M. analysed WGBS and Nanopore data and performed imputation analysis. M.T., A.F. and S.B. interpreted the results. M.T., A.F. and S.B. wrote the manuscript.

## Availability of data and materials

All raw sequencing data and methylation calls files are freely available through the European Nucleotide Archive (accession no. XXX).

## Supplementary Figures

**Supplementary Figure 1.**
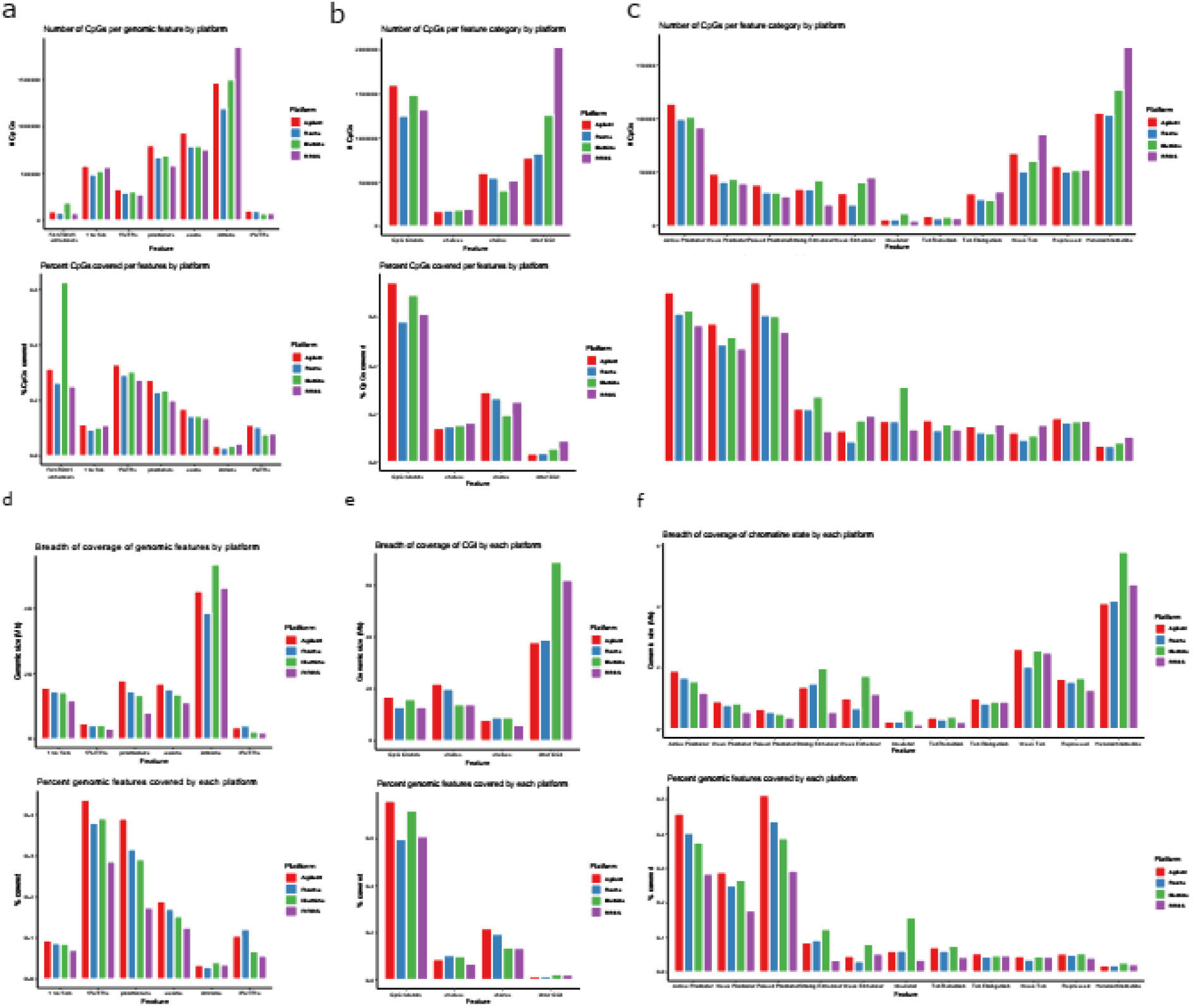
Feature annotation of region and CpGs targeted by design per platform. **(a)** Number of CpGs (top) targeted by design stratified by genic feature type and percent of all CpGs (bottom) in that feature type for each platform. **(b)** Number of CpGs (top) targeted by design stratified by CpG density feature type and percent of all CpGs (bottom) in that feature type for each platform. **(c)** Number of CpGs (top) targeted by design stratified by regulatory feature type and percent of all CpGs (bottom) in that feature type for each platform. **(d)** Size of the region in Mb (top) targeted by design stratified by genic feature type and percent of the total size of that feature type (bottom) for each platform. **(e)** Size of the region in Mb (top) targeted by design stratified by CpG density feature type and percent of the total size of that feature type (bottom) for each platform. **(e)** Size of the region in Mb (top) targeted by design stratified by regulatory feature type and percent of the total size of that feature type (bottom) for each platform.

**Supplementary Figure 2.**
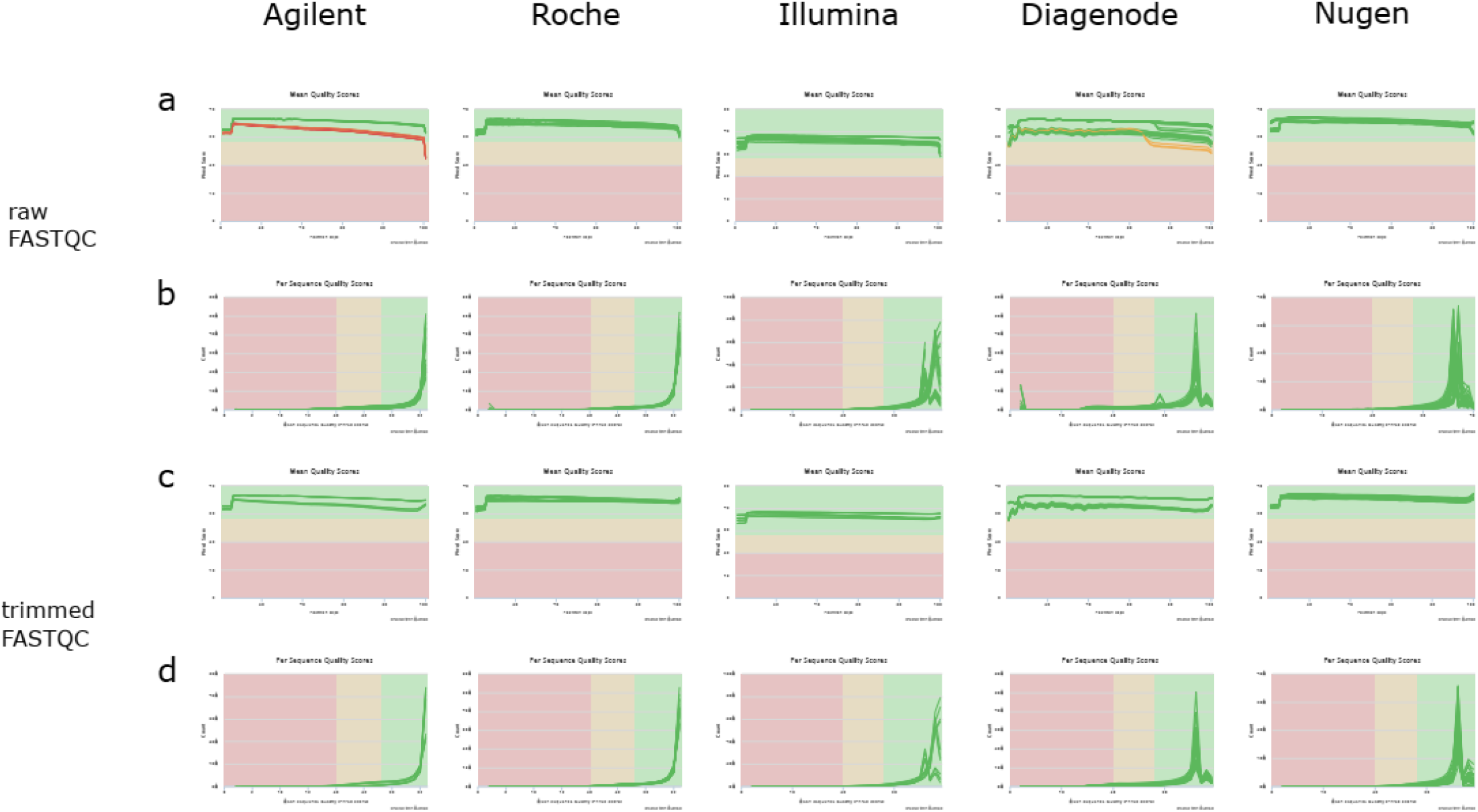
Raw and trimmed sequencing data quality by the platform. **(a)** Sequence quality (Phred score) by read position for raw FASTQ reads by the platform. **(b)** The number of raw reads as a function of sequence quality (Phred score) by the platform. **(c)** Sequence quality (Phred score) by read position for trimmed FASTQ reads by the platform. **(d)** The number of trimmed reads as a function of sequence quality (Phred score) by the platform.

**Supplementary Figure 3.**
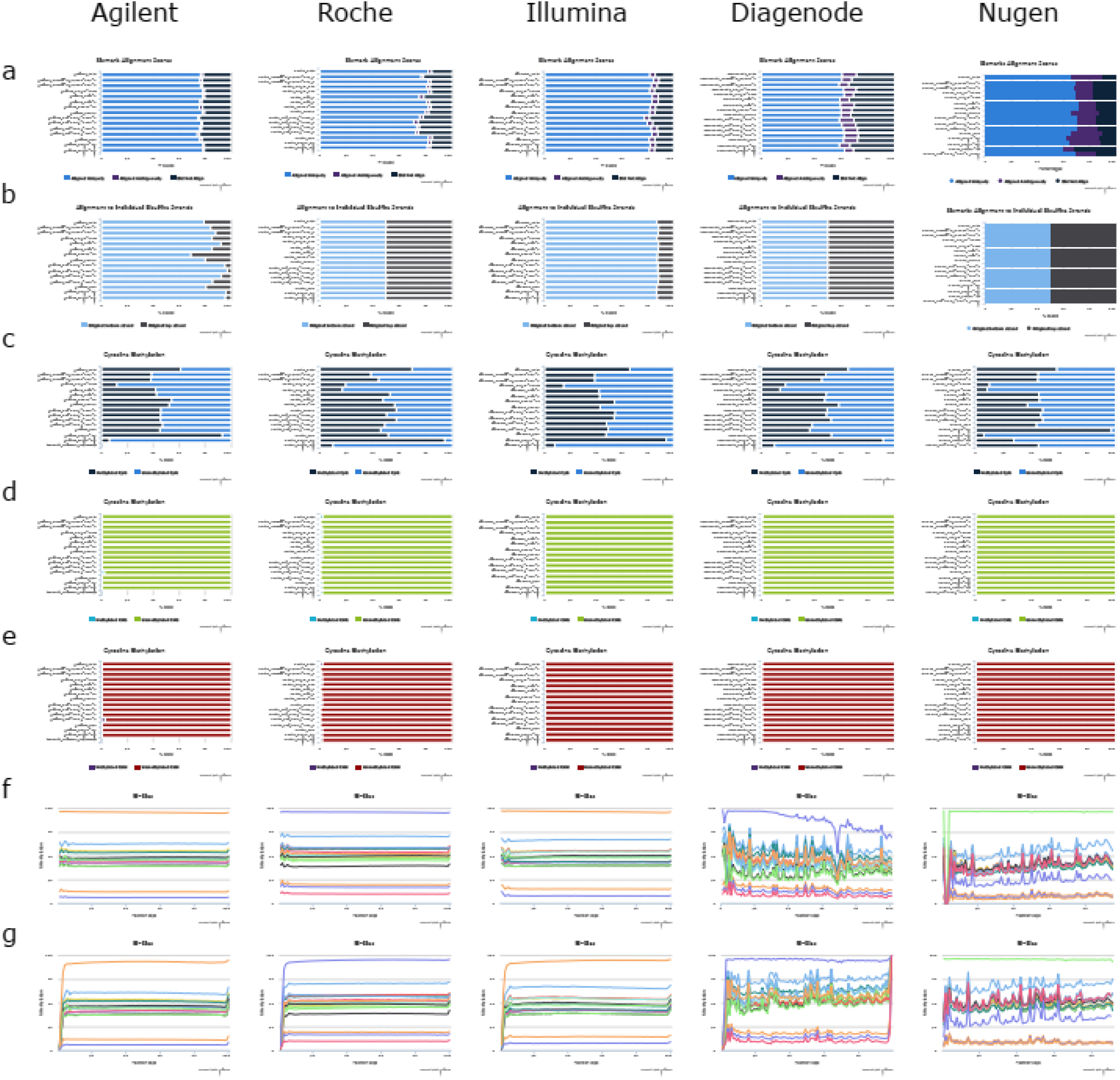
Sequencing data processing quality metrics. **(a)** Bismark alignment rates for uniquely, ambiguously, or unaligned reads for each sample by platform **(b)** Percent of reads aligning to top or bottom DNA strand for each sample by the platform. **(c)** Global methylation levels of CpG dinucleotides for each sample by the platform. **(d)** The global cytosine methylation level in CHG context for each sample by the platform used an estimate of sodium bisulfite under-conversion rates. **(e)** The global cytosine methylation level in CHH context for each sample by the platform used an estimate of sodium bisulfite under-conversion rates. **(f)** Methylation bias for the forward sequencing read. **(f)** Methylation bias for the reverse sequencing read

**Supplementary Figure 4.**
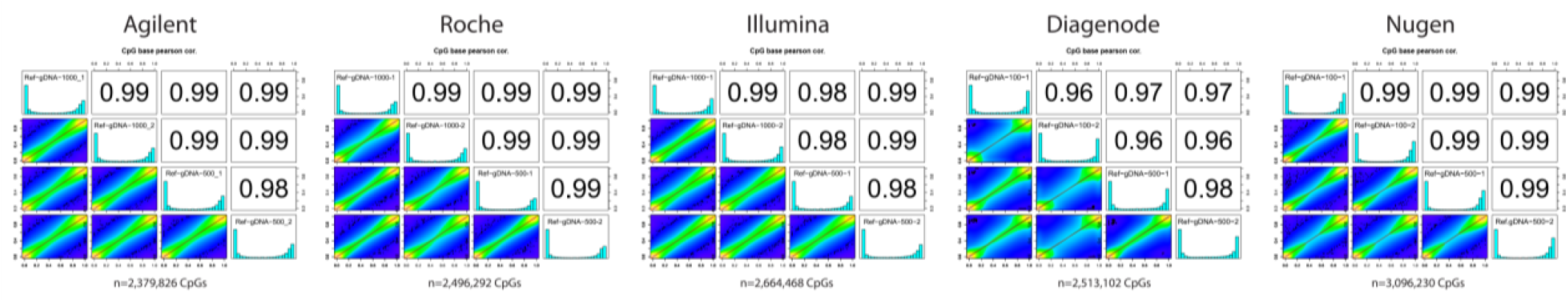
Concordance in DNA methylation levels dependent on sample input. Correlogram showing Pearson correlation coefficient of Ref.gDNA samples at specified inputs per platform.

**Supplementary Figure 5.**
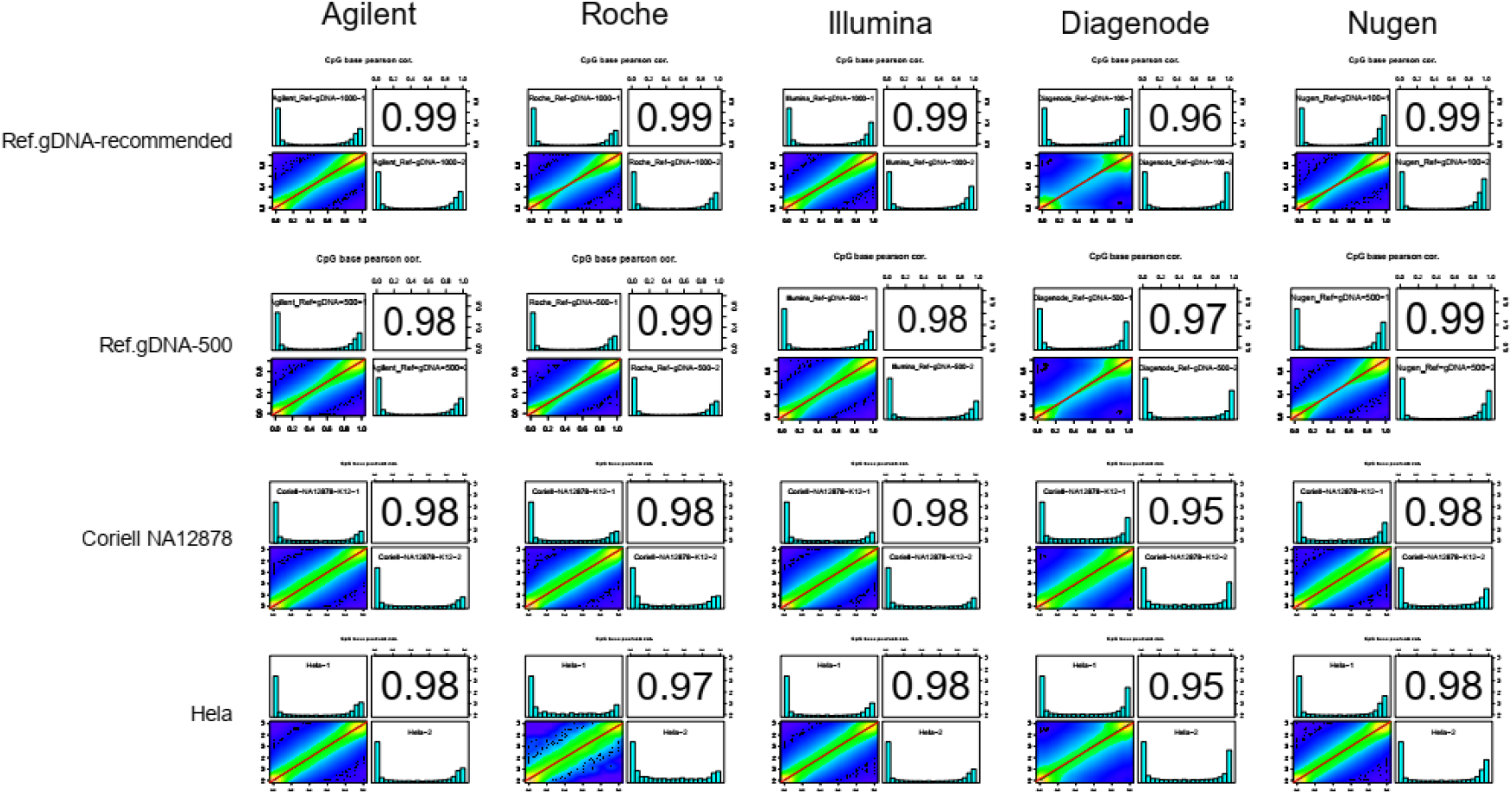
Correlogram showing intra-platform reproducibility in DNA methylation calls as pairwise Pearson correlation coefficient for each technical replicate sample by the platform.

**Supplementary Figure 6.**
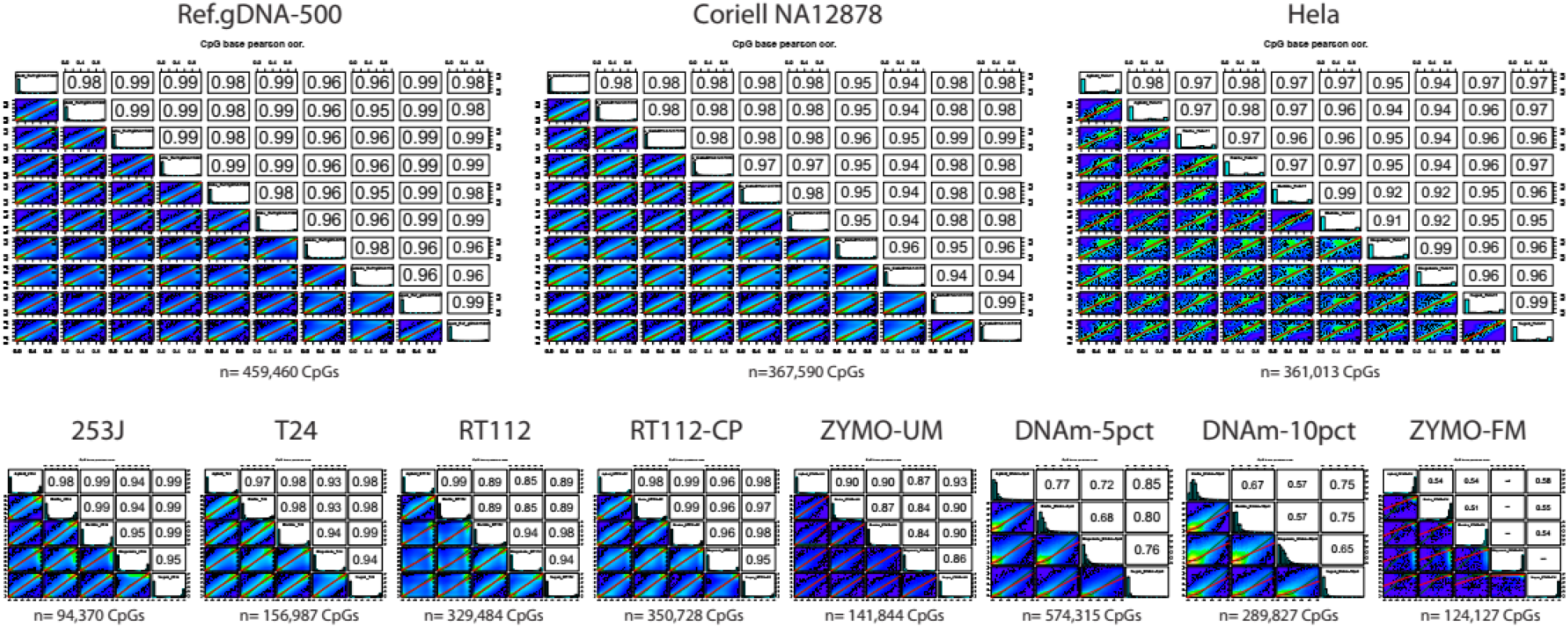
Correlogram showing inter-platform concordance in DNA methylation calls as pairwise Pearson correlation coefficient and the number of commonly interrogated CpGs at 10x for each sample across all platforms.

**Supplementary Figure 7.**
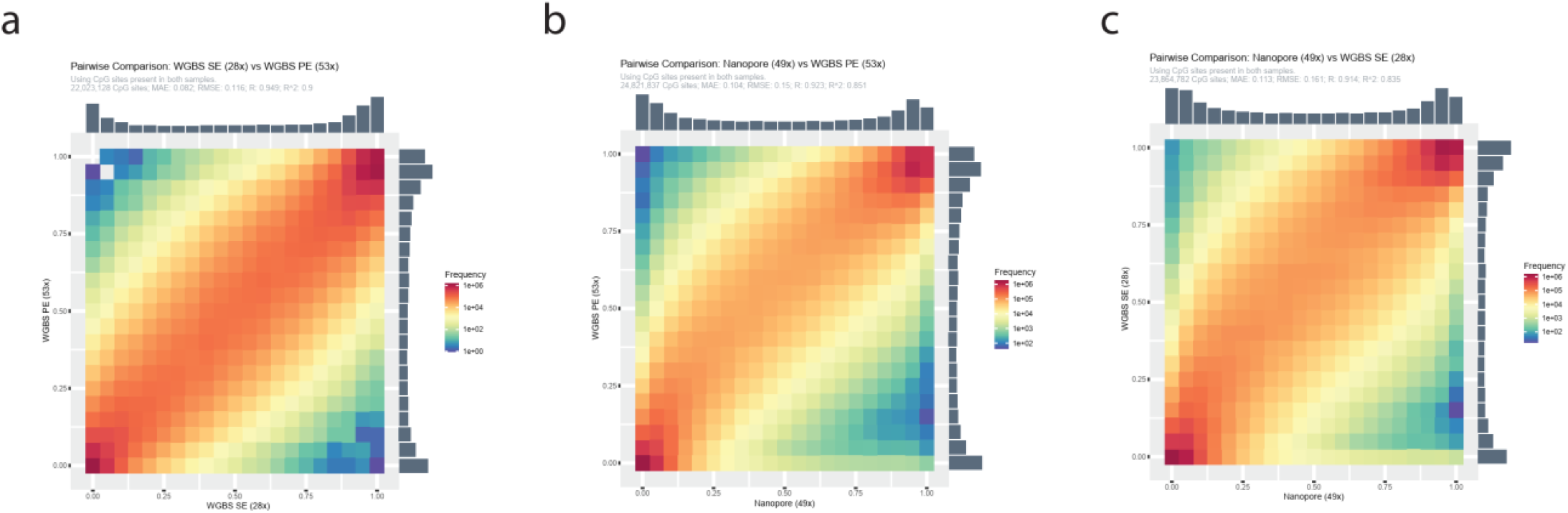
Scatterplot showing inter-platform concordance in DNA methylation calls as pairwise Pearson correlation coefficient for Coriell NA12878 data from **a)** WGBS SE vs PE **b)** Nanopore vs. WGBS PE, and **c)** Nanopore vs. WGBS SE

**Supplementary Figure 8.**
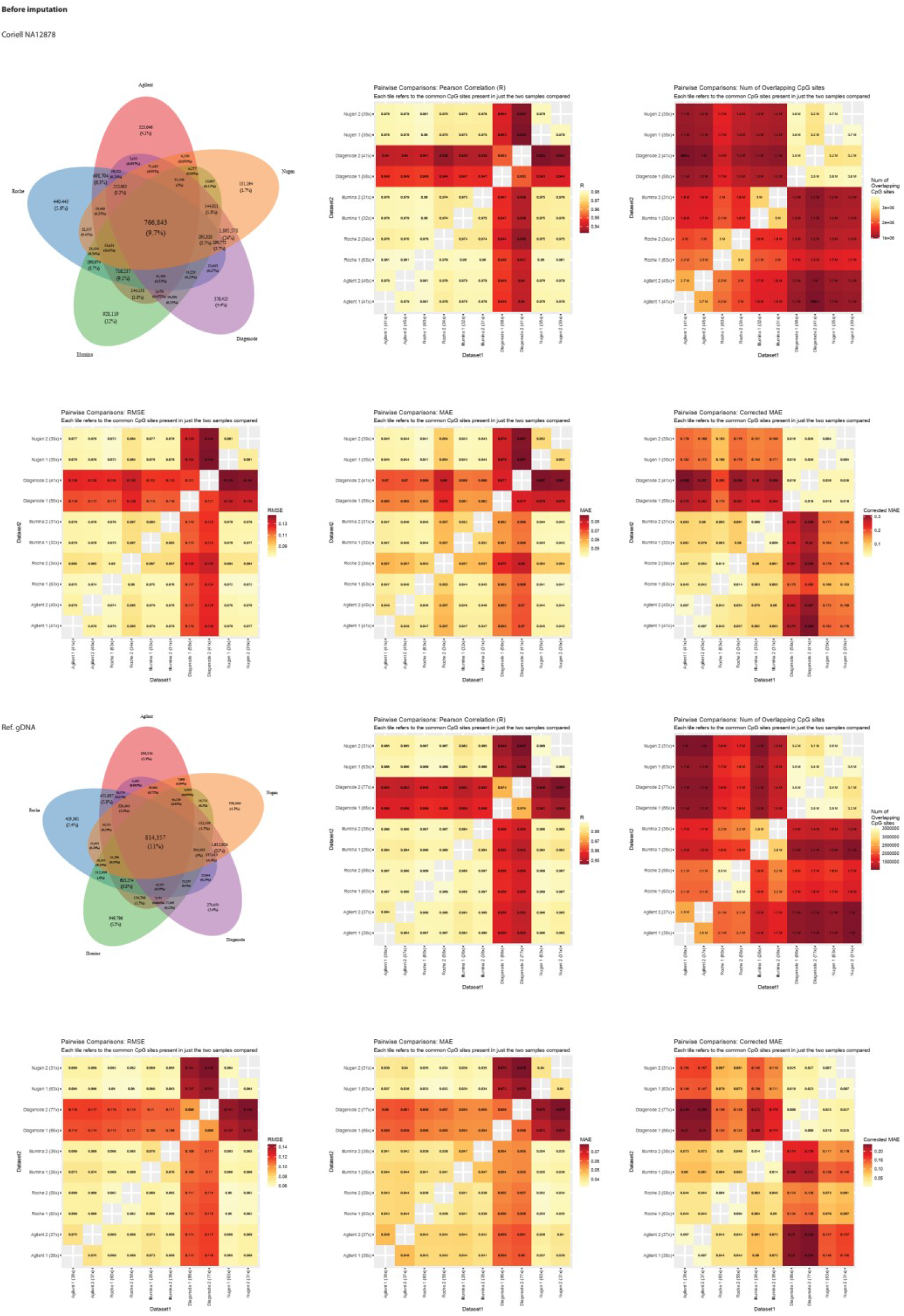
Interoperability between platforms before imputation for **a)** Coriell NA12878 and **b)** Ref.gDNA. Venn diagram showing CpGs overlapping between the platforms (top left), pairwise Pearson correlation coefficient (top centre), number of overlapping CpGs (top right), round mean squared error (RMSE) (bottom left), mean absolute error (MAE) (bottom centre) and corrected MAE for the number of sites present (bottom right).

**Supplementary Figure 9.**
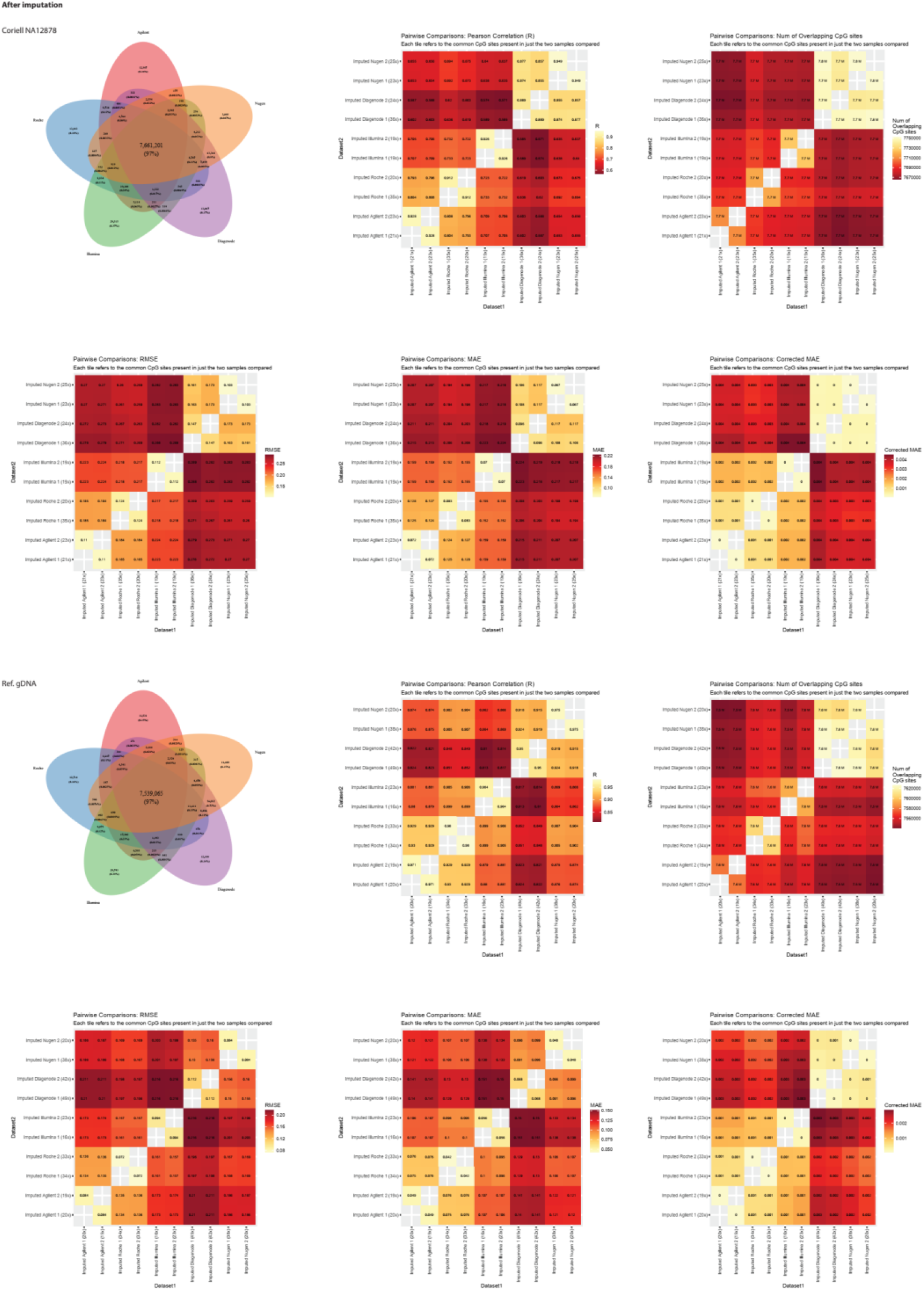
Interoperability between platforms after imputation for **a)** Coriell NA12878 and **b)** Ref.gDNA. Venn diagram showing CpGs overlapping between the platforms (top left), pairwise Pearson correlation coefficient (top centre), number of overlapping CpGs (top right), round mean squared error (RMSE) (bottom left), mean absolute error (MAE) (bottom centre) and corrected MAE for the number of sites present (bottom right).

**Supplementary Figure 10.**
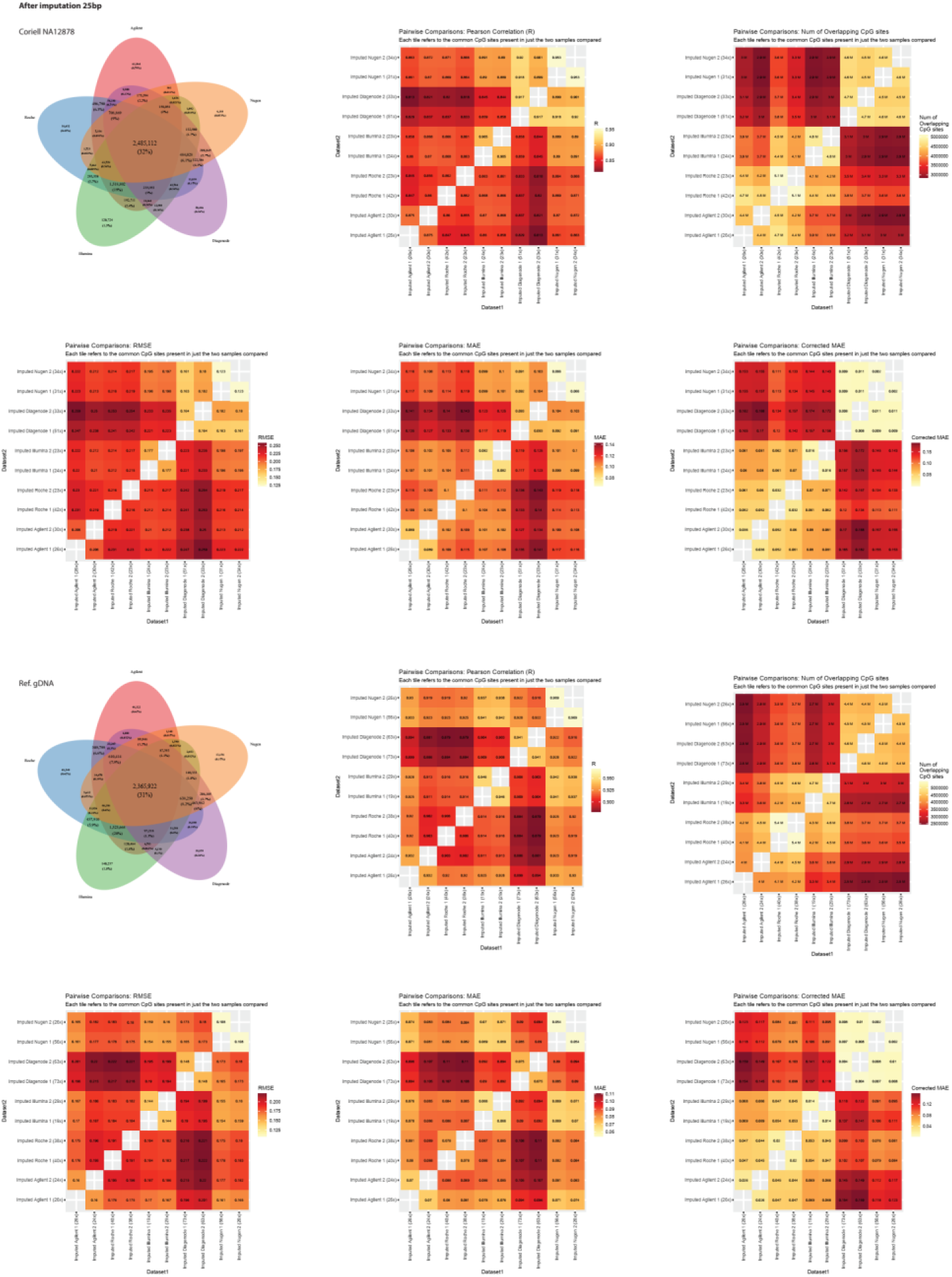
Interoperability between platforms after imputation with filtering for adjacent CpGs < 25bp for **a)** Coriell NA12878 and **b)** Ref.gDNA. Venn diagram showing CpGs overlapping between the platforms (top left), pairwise Pearson correlation coefficient (top centre), number of overlapping CpGs (top right), round mean squared error (RMSE) (bottom left), mean absolute error (MAE) (bottom centre) and corrected MAE for the number of sites present (bottom right).

